# Computational pipeline for targeted integration and variable payload expression for bacteriophage engineering

**DOI:** 10.1101/2024.06.17.597714

**Authors:** Jonas Fernbach, Emese Hegedis, Martin J. Loessner, Samuel Kilcher

## Abstract

Bacteriophages offer a promising alternative to conventional antimicrobial treatments, particularly in cases where such treatments have proven ineffective. While naturally occurring phages serve as viable options for phage therapy, advances in synthetic biology and genome engineering enable precise modifications to phages to enhance their therapeutic potential. One such approach is the introduction of antimicrobial genetic payloads into the phage genome. Conventional practice is to integrate such payloads behind genes expressed at very high levels late in the infection cycle, such as the major capsid gene (*cps*). Nevertheless, phages engineered to contain antimicrobial payloads are often difficult to obtain. For instance, the high expression of toxic payloads can prematurely halt host metabolism, leading to the failure in assembling viable phage progeny. To potentially expand the range of genes viable as genetic payloads, we developed a method to identify intergenic loci with favorable expression levels. We used the machine learning (ML)-based promoter prediction algorithm PhagePromoter to identify these loci. We then used this information to design a computationally-assisted engineering pipeline for the integration of genomic payloads at these locations. We validated our approach experimentally, engineering phages with bioluminescent reporter payloads at various predicted loci. We used the well characterized, strictly lytic, *Staphylococcus aureus* -infecting bacteriophage, K, as an engineering scaffold and employed homologous recombination to engineer three recombinant phages containing the reporter payload at different predicted loci throughout the genome. The recombinant phages exhibited expression levels consistent with our computational predictions and showed temporal expression patterns corresponding to their genomic locations in early, middle, or late gene clusters. Our study underscores the potential of integrating computational tools with classical sequence analysis to streamline the phage engineering process. This approach not only facilitates the rational design of phages with targeted payload insertions but also paves the way for high-throughput, automated phage engineering, fostering a new era of personalized phage therapy.

## Introduction

Bacteriophages, ubiquitous viruses with the innate capacity to infect and lyse bacteria, have long been considered potential alternatives to antibiotics. While antibiotics have largely supplanted bacteriophages in Western medicine, the escalating global prevalence of multi-drug resistant bacterial infections necessitates urgent exploration of alternative treatments like phage therapy [1, 2, 3, 4, 5]. Although numerous case studies demonstrate the efficacy of phages in treating antibiotic-resistant infections [6, 7, 8], natural phages present challenges such as the rapid emergence of phage-resistant bacteria [6, 9, 10], immunogenicity during extended treatment [4], and a restricted host range that can limit their applicability. To mitigate these challenges and augment the clinical utility of phages, various genetic engineering techniques have been devised to enhance their therapeutic efficacy [11, 12]. Specifically, the genetic ’arming’ of bacteriophages with antimicrobial effector payloads holds significant promise for advancing phage engineering in therapeutic contexts [12, 13]. A prevalent method for the precise, markerless integration of genetic payloads involves homologous recombination, which is commonly followed by either CRISPR-Cas9-assisted counterselection or plaque screening to identify recombinant phages [11]. The conventional approach places payloads downstream of structural genes that are highly expressed late in the infection cycle. Two such genes are the major capsid gene (*cps*) and holin/endolysin (*ply*) cassette [14, 15]. Although this method has proven useful in the past [16, 17, 13, 18], several caveats exist and could potentially be mitigated through a more diverse selection of insertion loci. Inserting payloads after the *cps* gene can inadvertently affect transcription and may not consistently yield viable phage progeny—a concern applicable to all insertion sites, particularly those that are uncharacterized or inaccurately annotated. Furthermore, these locations restrict the temporal expression of the payload to the end of the infection cycle [14, 15]. In certain applications, early and rapid payload expression could be advantageous. For example, reporter phages, which express signaling payloads for pathogen detection in diagnostics, could benefit from swift payload expression for more rapid signal detection. Another imaginable approach is to increase payload effect through the rapid and high expression of intracellular-acting proteins early in the infection cycle. This enhancement could also bolster antimicrobial efficacy against bacterial hosts equipped with innate or adaptive defense mechanisms. Low or intermediate expression levels may also be better suited for payloads where phage amplification at the infection site is beneficial, but high toxicity of the expressed payload may lead to premature cell death and abortion of the infection cycle. Given these considerations, there is a need to explore insertion sites with variable expression levels throughout the infection cycle. Developing reliable methods for predicting these sites and their corresponding expression levels could significantly advance genome engineering and phage therapy.

Understanding the binding sites for transcription factors serves as an initial step in gene expression regulation and can be deduced from the genetic sequence. The growing sophistication of machine learning (ML) algorithms, particularly those that infer transcription factor and polymerase binding sites, coupled with the influx of new bacteriophage sequence data, supports the integration of this knowledge into phage engineering methodologies. This could enhance both the precision and flexibility of traditional phage engineering approaches. Here, we developed and assessed a tool that employs ML for promoter inference to predict payload expression at optimal insertion sites, integrating this approach into our existing phage-engineering pipeline. Utilizing homologous recombination (HR) and subsequent bioluminescent plaque screening, we engineered three *S. aureus* -targeting *Kayvirus* variants, each equipped with a bioluminescent nanoluciferase (*nluc*) reporter payload. These payloads were inserted at loci identified through our predictive pipeline. The well-studied biology of *Stapylococcus* phage K in the literature enabled us to align our computational predictions and experimental outcomes with pre-existing gene expression data and validated promoter regions [14]. The engineered phages constructed in this study exhibited expression levels consistent with our predictions. This was true both for the absolute quantity of the expressed payload and the timing of expression. These findings were further corroborated by previously characterized temporal gene expression data for K. In summary, our newly developed tool enables quick identification of gene insertion sites, thereby offering a means to fine-tune payload expression levels in therapeutic phage scaffolds by identifying and leveraging variable promoter strengths.

## Results

### Comparison of various promoter prediction algorithms

A variety of algorithms exist to date which are designed to predict promoters from genomic sequence data. These range from simple, consensus motif-based approaches [19] to more complex, state-of-the art ML algorithms such as support vector machines (SVM), random forests (RF), and convolution neural networks (CNN) [20]. Our approach is designed to be universally applicable to whole-genome sequences, irrespective of phage or host species. This universality required that our chosen method for promoter prediction meet several pre-defined parameters. To allow for high-throughput processing of multiple genomes simultaneously, the method of choice had to be applicable to complete genome sequences and compatible with, e.g., a multi-fasta input format. Our approach is driven by the hypothesis that ML-based probability scores are inherently linked to promoter strengths. We therefore required machine-learning implementations which quantify the probability that a predicted promoter was correctly determined as such. A comprehensive overview of the algorithms evaluated for integration into our pipeline is presented in **Table 1**.

**Table 1:** Promoter prediction software and various parameters relevant for seamless integration into our pipeline.

| Tool | Multi-fasta | Big files | Promoter Core | Score or Probability | Output Format | Additional Notes | Citation |
| --- | --- | --- | --- | --- | --- | --- | --- |
| Bprom | No | Yes | Yes | Yes | Text on screen |  | [21] |
| bTSS-finder | Yes | Yes | Yes | Yes | Text file | Sequence length limited to 500bp. Not executable on MacOS. | [22] |

**Table 1: Promoter prediction software and various parameters relevant for seamless integration into our pipeline. (Continued)**
| Tool | Multi-fasta | Big files | Promoter Core | Score or Probability | Output Format | Additional Notes | Citation |
| --- | --- | --- | --- | --- | --- | --- | --- |
| BacPP | No | No | No | Yes | Text on screen |  | [23] |
| Virtual Footprint | No | Yes | Yes | Yes | Text on screen | Requires .gb extension. Sequence length limited to 1000bp. | [24] |
| IBBP | No | Yes | No | Yes | Text file | Windows only | [25] |
| iPro70-FMWin | Yes | Yes | No | Yes | Text on screen |  | [26] |
| iPro70-PseZNC | Yes | Yes | No | Yes | Text on screen | Fasta input file requires .txt extension. | [27] |
| iPro54-PseKNC | Yes | Yes | No | Yes | Text on screen | Fasta input file requires .txt extension. | [28] |
| 70ProPred | Yes | Yes | No | No | Text on screen | No input file (has to be entered manually on webpage), | [29] |
| CNNProm | No | No | No | Yes | Text on screen |  | [30] |
| MULTiPly | Yes | Yes | No | No | Text on screen | Slow for large datasets | [31] |
| iPromoter-2L | No | No | No | No | Text on screen |  | [32] |
| Promoter-Hunter | No | No | Yes | Yes | Text on screen | Requires predefined motif matrix | [33] |
| SAPPHIRE | Yes | Yes | Yes | Yes | Text file |  | [34] |
| XSTREME | Yes | Yes | Yes | Yes | Text file |  | [35] |
| Phage-Promoter | Yes | Yes | Yes | Yes | Text file | Trained on validated phage-encoded promoters, *S. aureus* specific | [36] |

We detail each method’s specifications concerning our pre-determined parameters, as well as additional factors which ruled out different methods. Ultimately, we opted for the recently developed algorithm PhagePromoter, which employs both support vector machine (SVM) and artificial neural network (ANN) methodologies [36]. This algorithm was the only candidate that met all our predefined criteria. PhagePromoter has the added benefit of being adapted towards the prediction of phage-specific promoters based on experimentally verified, phage-specific input training data.

### Promoter prediction and determination of suitable genetic loci for payload insertion in *S. aureus* phage K

We chose the well-characterized, strictly lytic *S. aureus* -infecting *Kayvirus* K as our engineering scaffold [37]. The annotated, complete sequence record was used (Genbank: KF766114.1).To maintain simplicity in our approach, we primarily concentrated our promoter search within intergenic regions. Acknowledging the potential for promoter regions to overlap with coding sequences, we expanded our search to include 50 base pairs at both ends of each intergenic region. The gene product *gp141* of K, annotated as comprising two exons interspersed with genes *gp142* and *gp143*, was considered as a single contiguous sequence to eliminate potential insertion sites between the exons. We utilized the Phage-Promoter Galaxy Docker build [38] for promoter predictions in intergenic regions. This approach revealed 303 promoters, with many achieving the maximum predictive score of 1 **(Figure 1A)**. We classified regions of high promoter element density within the intergenic spaces—termed Intergenic Promoter Regions (IPR)—and formulated a scoring algorithm to predict downstream gene expression levels. Individual PhagePromoter scores were exponentially weighted to place more emphasis on high-scoring predictions **(Figure 1B)**. The IPR score was calculated as the cumulative, weighted scores of all predicted promoters within that IPR. Previous studies have characterized the transcription landscape for K using quantitative mRNA sequencing data [14]. This allowed us to compare our computational predictions to experimental promoter data and gain insight into the validity of our approach. Significant overlap between the experimental data and computational promoter predictions was evident **(Figure 2A)**. Our method included 71 of 83 of the promoters experimentally validated in [14]. As posited in our initial hypothesis, the temporal variations in gene expression during an infection cycle could potentially influence the efficacy of specific payloads. The availability of temporal expression data for K [14] **(Figure 2A and B)** motivated us to select high-scoring IPRs for genomic regions expressed at different time points during infection. Genes expressed early, late or at an intermediate time during the infection cycle were determined using available data and the highest-scoring IPRs were identified for each temporal category. Only loci with a proximal downstream gene in the same orientation as the IPR were considered. Insertion sites were selected immediately downstream of the first gene following an IPR. This resulted in 3 IPRs at nucleotide positions 229-518 (early gene), 37505-38806 (late gene) and 119132-119495 (middle gene) **(Figure 2B)**.

**Figure 1.**
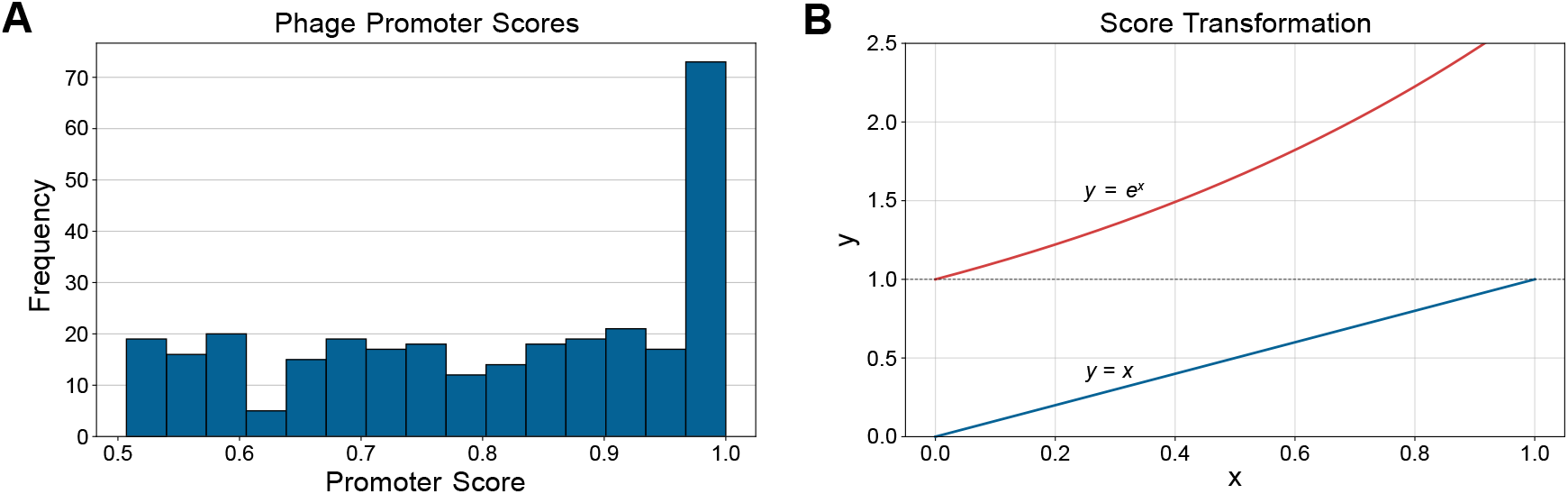
Promoter score distribution and weighting scheme. (**A**) Histogram of all predicted promoter scores across the phage K genome. **(B)** Original predicted promoter scores (blue line) were exponentially weighted to favor high scoring promoters (orange line).

**Figure 2.**
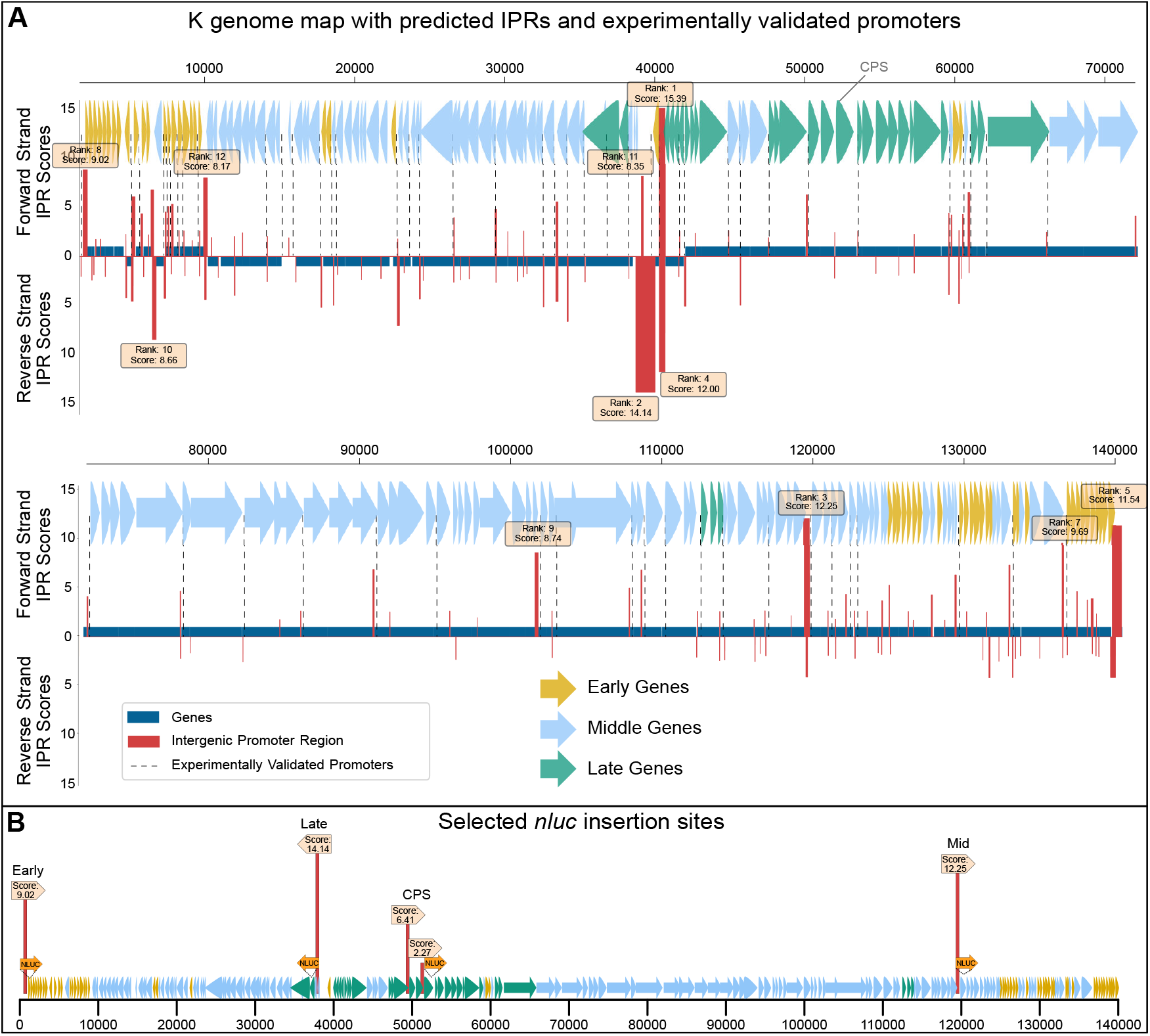
Genome map of phage K, including promoter predictions and selected insertion sites. **(A)** K genome map with predicted intergenic promoter regions (IPRs) and computed promoter scores. Genes are colored based on previously determined temporal mRNA expression and experimentally validated promoters are indicated with vertical dotted lines (adapted from [14]). All calculated IPRs are shown as vertical red bars (weighted scores for the 12 highest scoring IPRs are depicted above/below). **(B)** Reporter gene (*nluc*) insertion sites were selected based on the highest scoring IPRs which had a downstream gene behind which insertion of the payload was possible. IPRs with no immediate downstream gene were not considered. One insertion site each was selected for early, middle and late expressed genes. The CPS insertion site used in a previous study [18] is indicated.

### Engineered K::*nluc* variants show bioluminescence expression corresponding to predicted promoter strengths and temporal gene expression data

To engineer the three reporter phages that encode *nluc* at different IPRs (early, middle, late), we used an HR-mediated approach. To this end, we constructed homology donor plasmids (pEDITnluc_early_, pEDITnluc_middle_, pEDITnluc_late_) that guide site-specific recombination (**Figure 3A**). HR was performed during infection of the K-susceptible host RN4220 (**Figure 3B**). The resulting phage lysates contained a mixture of wildtype and recombinant phage. Recombinant phages were enriched by bioluminescence screening (**Figure 3B+C**), isolated, sequenced, and purified by CsCl ultracentrifugation to remove residual NLuc protein from the phage preparation. To quantify NLuc expression levels and kinetics, bioluminescence measurements were performed over a timespan of 140 min at 37°C **(Figure 4A)**. The observed expression levels aligned with our computational predictions. Specifically, the phage variant K::*nluc*_Late_ exhibited the highest bioluminescence, registering an IPR score of 14.14, followed by K::*nluc*_Mid_ (IPR score: 12.25) and K::*nluc*_Early_ (IPR score: 9.02). We additionally measured bioluminescence of the previously engineered K::*nluc*_CPS_ [18], which has the *nluc* gene inserted behind the major capsid protein coding gene (*cps*). The bioluminescence of K::*nluc*_CPS_ was observed to be approximately 5 times higher than that of K::*nluc*_Late_, despite the prediction of only a single, short promoter element (ATAAAT, PhagePromoter score: 0.921, IPR score: 2.51) upstream of *cps*. Interestingly, an IPR with a higher score (IPR score: 6.406) was identified two genes upstream (1832 bp) of *cps*. This IPR coincides with a location of an experimentally validated promoter and could potentially be contributing to the elevated transcription levels observed (**Figure 2A**). The latency period preceding the onset of expression was short for all phage variants, and all phages produced detectable bioluminescence within the first 10 minutes of infection. To more clearly delineate kinetic differences at the initial stage of infection, the same assay was conducted at 25°C for a duration of one hour (**Figure 4B**). K::*nluc*_Early_ showed the fastest increase in bioluminescence, followed by K::*nluc*_Mid_ and K::*nluc*_Late_ with similar profiles. K::*nluc*_CPS_ showed the longest time until onset of expression.

**Figure 3.**
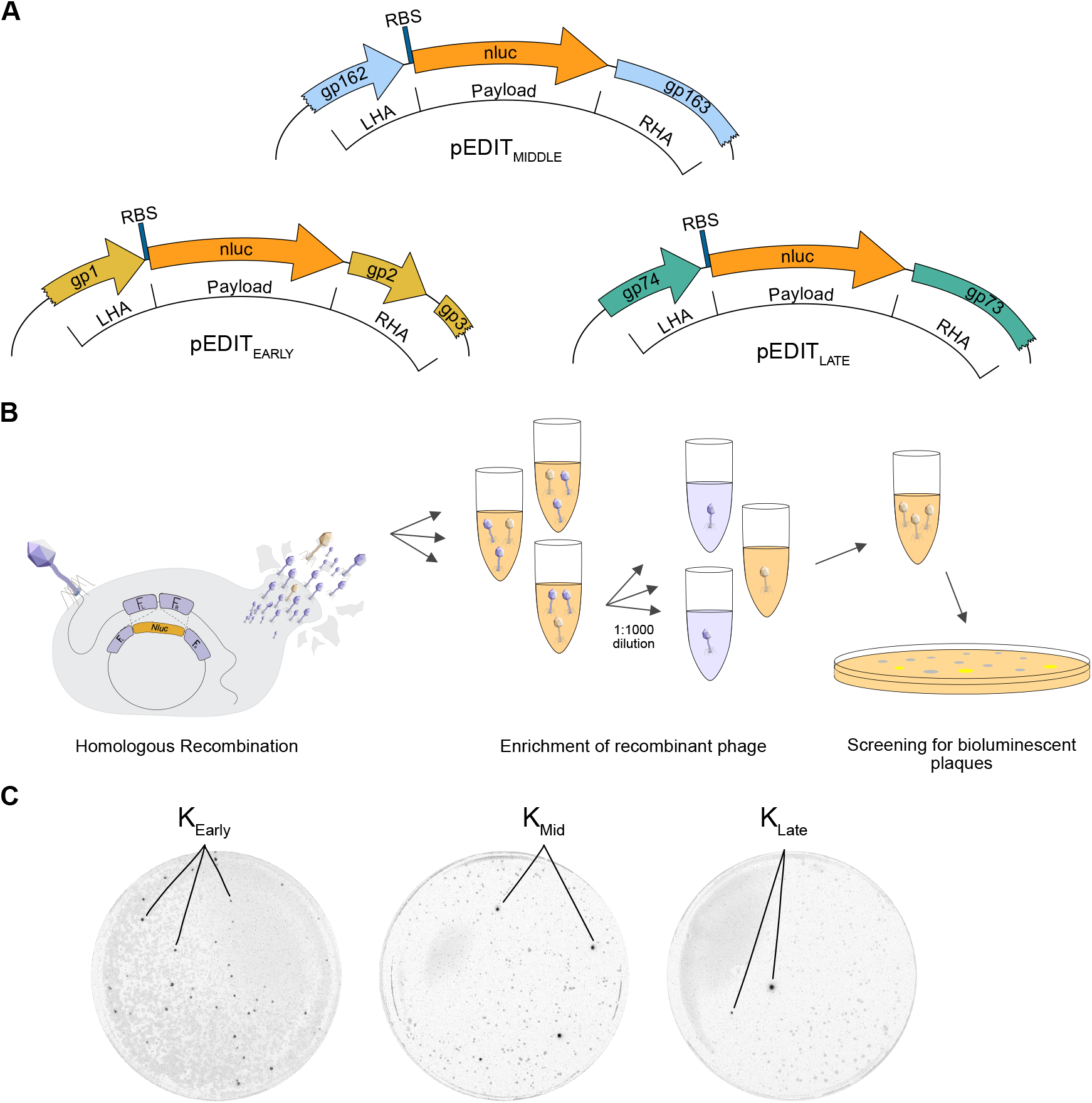
Bioluminescence screening for recombinant K::*nluc* phages. **(A)** Lysates obtained from HR were used to obtain escape mutants potentially containing the intended insertion. Lysates were diluted 1000-fold and evaluated for bioluminescence emission in liquid infection assays (1*10^4^ PFU/ml phages (initially), 1*10^6^ CFU/ml bacteria). For positive samples, this process was repeated two more times. Finally, plaque assays were performed and individual plaques showing bioluminescence isolated, purified and Sanger sequenced to validate correct insertion of the nanoluciferase gene. **(B)** Detection of bioluminescent plaques for phages K::*nluc*_Early_, K::*nluc*_Mid_ and K::*nluc*_Late_.

**Figure 4.**
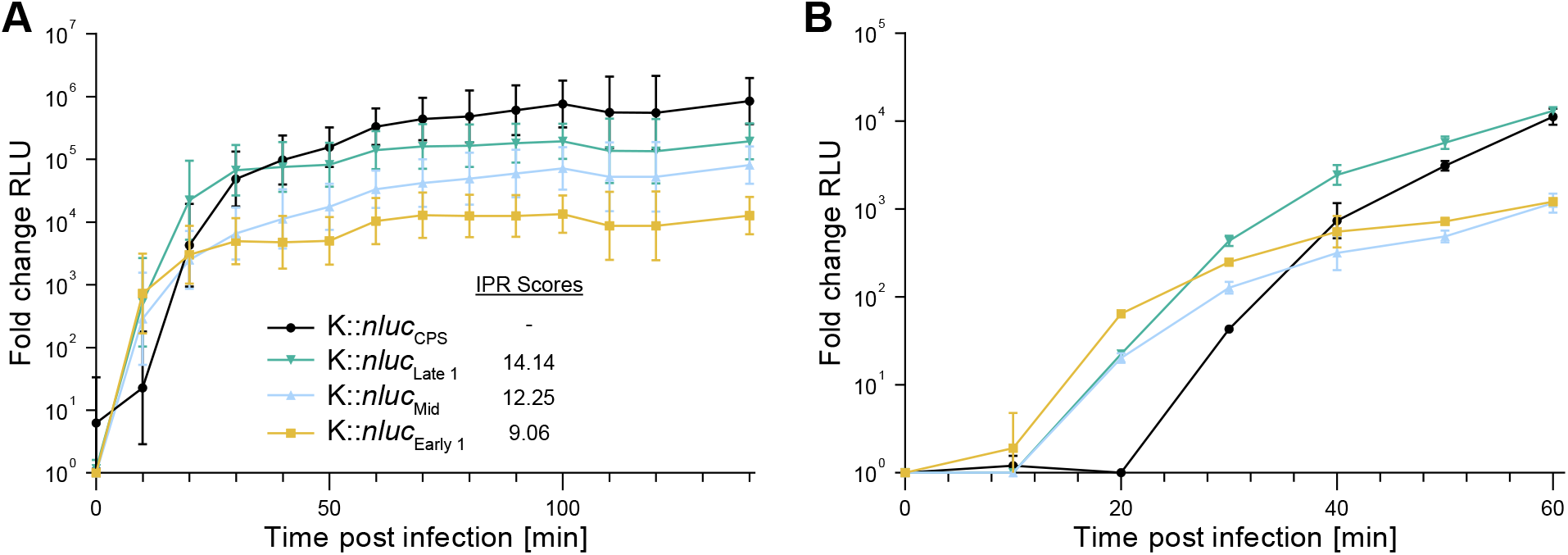
Bioluminescence time course measurements. **(A)** Bioluminescence time course measurements for K::*nluc*_Early_, K::*nluc*_Late_ and K::*nluc*_Mid_, as well as a previously characterized K::*nluc*_CPS_ [39] were obtained by calculating the fold change in relative light units (RLU) compared to infection with K wildtype at an initial bacterial density of OD_600_=0.01 and phage titer of 1*10^8^ PFU/ml. K propagation host *S. aureus* PSK was used as a bacterial host and infection occurred at 37°C for 140 min. Values are corrected for background luminescence. **(B)** Bioluminescence time course measurements for phages, bacteria and concentrations described in **(A)** were performed at 25°C for 1h to elucidate differences in temporal expression. Data is mean +/- standard deviation from biological triplicates.

## Discussion

Phage engineering offers a promising avenue for augmenting the therapeutic applications of naturally occurring phages. While the field has seen a surge in diverse genetic engineering techniques [11, 12, 40, 41], the challenge of selecting appropriate payload insertion sites within the phage genome remains underexplored. As mentioned previously, traditional approaches have largely relied on empirical methods, focusing on regions behind late-expressed structural genes such as the major capsid protein [16, 17, 13, 18]. Inspired by recent advancements in computational sequence analysis, including ML algorithms for promoter and transcription factor binding site identification, we aimed to explore and alternative integration sites.

Promoters serve as pivotal elements in the regulation of gene expression, acting as short sequence motifs typically located upstream of the genes they regulate. These sequences facilitate the binding of *σ*70-RNA polymerase (RNAP) and initiate transcription at the transcription start site [42]. Research indicates that initiation rates and RNAP kinetics are influenced by a number of factors such as sequence context, adjacent motifs, and the overall genomic landscape. These factors can lead to variations in the kinetics of RNA polymerase (RNAP), affecting both its recruitment to the promoter region and its processivity during transcription elongation. Some studies have reported differences of up to 10,000-fold in these kinetic properties, depending on the specific promoter region involved [43, 44, 45, 46, 47]. Promoters that closely align with the consensus sequence are generally associated with stronger RNAP affinity, a factor that can contribute to elevated transcription levels [47]. However, strong RNAP binding can sometimes lead to over-stabilization of the initiation complex at the transcription start site, resulting in lower levels of gene expression [48]. This phenomenon has been observed primarily in genes with high baseline expression levels and for small, single nucleotide sequence variations [48]. Recent studies have also challenged traditional assumptions about promoter orientation and transcription initiation. One study shows an inherent bi-directionality of promoters, putting into question the traditional assumptions of promoter orientation and transcription initiation [49]. Interestingly, we frequently observed the co-occurrence of predicted promoter regions on both strands, which is possibly a result of this promoter-based DNA-sequence symmetry (**Figure 2A**). Moreover, the genomic characteristics of specific bacterial species can introduce additional complexities. For example, the low G/C content in *Staphylococcus* species may lead to an over prediction of TATA motifs, characteristic of the -10 promoter region. Additionally, purine-pyrimidine base preferences can influence the energetic stability of the DNA double helix, thereby affecting transcription independent of promoter sequence motif conservation [46].

In the quest to identify optimal sites for genetic payload integration, we leveraged ML methodologies to augment traditional sequence analysis. Specifically, we utilized Phage-Promoter [36], a tool designed for predicting phage promoters, which we selected after a comprehensive evaluation of existing promoter prediction methods (see **Table 1**). This tool was particularly advantageous for our study for several reasons. Firstly, Phage-Promoter is trained on a large dataset comprising both bacterial and phage-encoded promoters, along with their corresponding gene expression levels. This extensive training data allows the algorithm to generate probability scores that serve as a proxy for promoter strength. This is in contrast to conventional, non-ML-based consensus methods, where the general distance to the consensus sequence of a promoter is often used as a metric. We hypothesized that the ML-generated probability scores could serve as a reliable indicator of promoter strength. However, our study has its limitations, primarily due to its inability to consider various complex factors influencing bacterial transcription regulation, such as transcription factors, RNA binding proteins, RNA secondary structure, RNAP regulators, intricate promoter architecture and late transcriptional regulators (Ltr) specific to phages infecting Gram-positive bacteria [50, 51, 52, 53, 54, 55, 56]. The inaccuracies in predicting the expression levels of certain genes, like *cps*, highlight these limitations and suggest the need for further investigation to refine our ML-based promoter prediction model.

To enhance the reliability of our predictions, we devised a scoring metric that weights individual promoter scores based on their probability. These weighted scores are then clustered into IPRs, providing a holistic approach aimed at reducing false positives. However, this method is still in its nascent stage and may exclude potentially viable insertion sites, constituting false negatives.

While we have benchmarked our approach using PhagePromoter, the pipeline is inherently modular, allowing for the integration of alternative or additional promoter prediction methods. This adaptability makes our pipeline a valuable asset for future high-throughput, automated phage engineering endeavors.

In this study, we leveraged the genetic manipulability of the well-established *S. aureus* phage K to integrate a bioluminescent nanoluciferase (*nluc*) reporter payload at selected loci, a strategy informed by previous successful applications of *nluc* as a reliable gene expression marker [17, 18, 57]. This facilitated the development of recombinant phages K::nluc_Early_, K::nluc_Mid_, and K::nluc_Late_, each embodying the highest-scoring insertion sites within early, middle, and late gene clusters, respectively. Our bioluminescence time-course measurements revealed distinct kinetic profiles for these phages, aligning well with the anticipated temporal expression patterns of their respective gene clusters [58, 14, 15]. These observations not only corroborate the predictions made by our ML-based promoter analysis but also underscore its potential as a robust tool in the rational design of phage engineering projects.

Our work highlights the synergistic potential of combining advanced ML algorithms with traditional sequence analysis to inform and enhance phage engineering strategies. The flexibility inherent in our modular pipeline facilitates the incorporation of diverse promoter prediction methods, allowing for adaptability and refinement in response to evolving analytical tools and techniques. We believe that the development of such integrative approaches is pivotal for advancing high-throughput, automated phage engineering, opening up possibilities for more accurate sequence-based expression predictions. This, in turn, empowers researchers to perform targeted payload insertions with heightened precision and reliability, paving the way for innovations in phage therapy and biotechnology.

## Materials and Methods

### Bacterial strains and culture conditions

*S. aureus* laboratory strain RN4220 (DSM 26309) was used as an engineering host to generate all K::*nluc* variants. PSK was used as propagation host for K and all K::*nluc* variants. *E. coli* XL1-blue MRF’ (Stratagene) was used for plasmid amplification prior to transformation of RN4220. *E. coli* were grown in Luria-Bertani (LB) liquid medium (3M NaCl, 10g/l tryptone, 5g/l yeast extract) overnight at 37°C (180 rpm). *Staphylococcus* species were grown in Brain Heart Infusion (BHI, Biolife Italiana) broth overnight at 37°C (180 rpm).

### DNA amplification

Reaction mixes consisted of 19 *µ*l H2O, 2.5 *µ*l of forward and reverse primer respectively, 1 *µ*l template and 25 *µ*l 2x Phusion High-Fidelity PCR Master mix with HF buffer (Thermo Fisher). The template amount was varied in cases where fragment amplification proved difficult. PCR reactions were conducted with the following conditions: 5 min at 95°C, 30 sec at 95°C, 30 sec at annealing temperature, (30 sec per 1000 bp) at 72°C, 5 min or 10 min at 72°C. Steps 2 to 4 were repeated for 30-35 cycles. The annealing temperature was varied in cases where fragment amplification proved difficult. Gel electrophoresis was performed in 1% agarose gel (1% agarose in 1x TAE buffer) at 110 V. Samples were loaded as follows: 9 *µ*l H2O, 2 *µ*l 6x DNA loading dye (Thermo Fisher) and 1 *µ*l PCR product. As DNA ladder, GeneRuler 1 kb DNA ladder was used. The loading dye was supplemented with 100x GelRed™ Nucleic Acid Gel Stains (Biotium).

### Plasmid construction

Design of homologous recombination (HR) plasmids were done analogous to [18]. The plasmid pLEB579 (kindly gifted by T. Takala, University of Helsinki, Finland) was used as a backbone. The pEDIT homology donor templates were constructed by integrating an *nluc* gene (optimized for *S. aureus* codon usage, avoiding of Rho-independent termination and an added upstream ribosome binding site: GAGGAGGTAAATATAT), flanked by homology arms corresponding to the intended K insertion site, into the linearized pLEB579 backbone. We chose the insertion site to be immediately downstream of the first gene following our predicted IPR. When the intergenic distance to the next gene (i.e., the second gene after the IPR) was below 20 base pairs (bp), we took measures to avoid disrupting the ribosomal binding site (RBS) of the downstream gene by artificially extending the intergenic region. This involved duplicating the required bases from the end of the preceding gene and inserting them right after the payload, thus preserving the integrity of the subsequent intergenic region. The homology arms were designed to be 200-400 bp in length and where possible to exclude full-length genes of unknown function to mitigate production of potentially toxic gene products during cloning. All synthetic sequences were acquired as GeneArt String DNA Fragments (Thermo Fisher), albeit the spacer sequences for pSELECT, which were ordered as GeneArt Gene Synthesis (Thermo Fisher). Strings and plasmid backbones were amplified by PCR followed by purification with the Wizard SV Gel and PCR Clean-up System (Promega). Plasmids were assembled using isothermal GibsonAssembly^®^ reaction (NEBuilder^®^ HiFi DNA Assembly Master Mix).

### Transformation of *E. coli* cells

To prepare electrocompetent *E. coli*, LB medium was inoculated with an overnight culture and grown until an OD_600_ of 0.4-0.6 was reached. The cells were incubated on ice for 30 min, centrifuged (4000 g, 15 min, 4°C) and pellets resuspended in 10 % glycerol. This was repeated three times and cells were stored at -80°C if not used directly. A maximum of 10 *µ*l of the DNA samples were transferred onto MF-Millipore 25 *µ*m MCE Membrane Filters (SigmaAldrich) in H2O and removed again after 10-20 min. Electrocompetent cells were mixed with the DNA and electroporated at 2.5 kV, 200 Ω, 25 *µ*F. 1 ml SOC medium (2% (w/v) Tryptone, 0.5% (w/v) Yeast extract, 10 mM NaCl, 2.5 mM KCl, 10 mM MgCl2, 20 mM Glucose) was added immediately and the cells in SOC were left at 37°C for 15 min without agitation, then with agitation (300 rpm) for 1h. Cells were grown overnight on selective medium at 37°C and successful transformants isolated.

### Transformation of *S. aureus* cells

To generate electrocompetent cells, BHI medium was inoculated with *S. aureus* and grown until an OD_600_=1 was reached. Cells were left on ice for 15 min, then washed three times with cold H2O and two times with 10 % glycerol (3000-5000 g, 10 min, 4°C). The cells were stored at -80°C if not used directly. For electroporation, electrocompetent cells were thawed for 5 min on ice and incubated for another 5 min at RT. They were centrifuged (5000 g, 5 min) and the pellet was resuspended in 70 *µ*l electroporation buffer (10 % glycerol, 0.5 M sucrose in H2O). DNA samples were transferred onto MF-Millipore 25 *µ*m MCE Membrane Filters (Sigma-Aldrich) in H2O and removed again after 10-20 min. DNA was then added to the cells and they were electroporated at 2.1 kV, 100 Ω, 25 *µ*F. 1 ml B2 medium (10 g/l casein hydrolysate, 5 g/l D-glucose, 1 g/l potassium phosphate dibasic, 25 g/l NaCl, 25 g/l yeast extract) was added immediately and the cells in B2 were left at 37°C for 2 h with agitation (300 rpm). The cells were then plated onto B2 plates supplemented with selective antibiotics and stored overnight at 37°C. Successful transformants were isolated the following day.

### Plasmid sequencing and evaluation

Sanger sequencing of PCR products (purified with Wizard SV Gel and PCR Cleanup System (Promega)) or plasmids (purified with GenELuteTM Plasmid Miniprep Kit (Sigma-Aldrich) was conducted at at Microsynth AG (Switzerland). Results were analyzed with CLC Genomics Workbench.

### Phage handling and storage

Phages were stored in S/M buffer (5.8 g/l NaCl, 2 g/l MgSO4, 6 g/l Tris-HCl (pH 7.5)) at 4°C. To pick individual plaques from plates, wide bore pipette tips were used to transfer a small amount of the plaque-containing soft-agar into 100 *µ*l S/M buffer.

### Spot-on-lawn and full plate overlay assays

To perform spot-on-lawn assays, 200 *µ*l of stationary phase bacterial culture were added to 5 ml molten (47°C) LC soft agar (10 g/l tryptone, 7.5 g/l NaCl, 1% D-glucose, 2 mM MgSO4, 10 mM CaCl2, 5 g/l yeast extract, 4 g/l agar), vortexed briefly and poured onto a 0.5x BHI plate. The soft agar was left to dry for at least 20 min at RT. 10 *µ*l spots of 10-fold decreasing phage dilutions were placed onto the agar. The spots were left to dry for 1 h and then stored overnight at 37°C. Full plate overlay assays were performed in a similar manner, with the difference that for each phage dilution, 10*µ*l were added to 5 ml molten LC soft agar together with the host bacteria and then poured onto a 0.5x BHI agar plate. The bacteria were added first, the LC soft agar tube was vortexed briefly, then the phages were added and transferred to the plate. The plates were dried for at least 30 min and incubated overnight at 37°C.

### Phage propagation and purification

To obtain a pure phage lysate, full overlays were performed with the phage on its host bacterium to obtain 3-6 plates with semi-confluent lysis. On the following day, 5 ml S/M buffer were added to each plate and placed at 4°C for 2-3 h with gentle agitation. The lysate was then collected and centrifuged (10’000 g, 10 min, 4°C) to pellet cell debris. The supernatant was sterilized by filtration using 0.22 *µ*m filters. Open-Top Thinwall Ultra-Clear Tube (Beckman Coulter) were loaded with the following CsCl density layers: 1.7 g/ml CsCl, 1.5 g/ml CsCl, 1.35 g/ml CsCl. The densities were obtained by dissolving CsCl salt in S/M buffer. As the topmost layer, the phage lysate sample adjusted to a density of 1.15 g/ml was added. Ultracentrifugation was performed at 20’700 rpm for 2 h at 10°C. The tubes were then collected and the visible phage band was recovered. The lysate was dialyzed for a minimum of 2x 2 h using the Slide-A-LyzerTM Mini Dialysis Device (Thermo Scientific).

### Engineering of K::*nluc*

The engineering process of K::*nluc* through HR was conducted analogously to the methods outlined in [18]. Briefly, the corresponding pEDIT plasmid for each engineered phage was transformed into *S. aureus* RN4220, creating the engineering strains RN4220 (pEDIT). These strains were then infected with serial dilutions of phage K using a softagar overlay method, with initial concentrations of 1 ∗ 10^4^ PFU/ml phages and 1 ∗ 10^6^ CFU/ml bacteria, to yield high titer lysates, obtained by washing semi-confluent plates with S/M buffer as previously described. Subsequent bioluminescence assays were conducted on the lysates using *S. aureus* PSK as the host. Lysates were diluted 1000-fold and evaluated for bioluminescence emission in liquid infection assays. Positive samples underwent two additional rounds of this dilution and assessment process. This assay was iteratively performed until the detection of single, bioluminescent plaques on full-plate overlays was possible. Finally, individual bioluminescent plaques were isolated, purified, and Sanger sequenced to confirm the correct and precise insertion of the nanoluciferase gene.

### Bioluminescence assays

For expression analysis of the purified recombinant phages, stationary phase bacterial cultures were diluted to OD_600_ 0.01, inoculated with 1∗10^8^ PFU/ml phage and incubated at 37°C (180 rpm agitation). Bioluminescence measurements were taken by combining 25 *µ*l of the sample solution with an equal volume of prepared buffer-reconstituted nluc substrate as detailed by the manufacturer (Nano-Glo Luciferase Assay System; Promega). Measurements were taken every 10 min in Nunc™ F96 MicroWell™ 446 plates using a GloMax^®^ navigator luminometer (Promega) with 5s integration time and 2s delay. To determine the background-corrected fold-change in relative light units (RLU) for each phage, measurements were normalized to a control reaction of each phage in BHI medium and the fold-change was calculated as the difference in RLU to an infection of the same strain with wildtype K. All measurements were performed in triplicate. To determine plaques containing recombinant, nanoluciferase-expressing phage from full plate overlays, 500 *µ*l of Nano-GloR substrate were spread onto the plate. Plates were photographed using Gel Doc XR+ Gel Documentation System, once with no illumination and 50 s exposure time, and once again using trans-white illumination and exposure of 0.2 s. Images were overlayed to determine the location of plaques showing bioluminescence on the full plate. For those engineered phages where no bioluminescent plaques were detectable, enrichment steps were conducted by liquid infection of PSK at an initial MOI of 0.01 (1 ∗ 10^4^ PFU/ml phages, 1 ∗ 10^6^ CFU/ml bacteria) until a significant rise in bioluminescence (*>* 10^3^ RLU) was detectable. The solution was then diluted 1:1000-fold and enrichment repeated. This was continued until bioluminescent plaques were detectable in the plaque assay.

### Bioinformatics

Evaluation of the various promoter prediction programs was conducted with the software versions as they were online available in the time between 02.06.2022 and 08.06.2022. The full genome sequence and annotations of K were acquired from genbank file KF766114.1. The PhagePromoter Galaxy Docker Build (Galaxy Version 0.1.0) was used for generating promoter predictions using the following parameters: Search both strands yes, threshold 0.5, phage family *Siphoviridae*, host bacteria genus *Staphylococcus aureus*, phage type virulent. Programming, plotting and sequence analyses were done in Jupyter notebook version 6.1.4 running python 3.8.5. CLC Genomics Workbench version 20.0.4 was used for additional sequence analyses such as primer and string design as well as evaluation of sanger sequencing results. OpenAI’s ChatGPT 4 [59] was used as a tool to assist with formatting and editing the manuscript. This involved iterative refinements to ensure clarity and conciseness of the content presented.

