## Supplemental Table 1 for "Computational pipeline for targeted integration and variable payload expression for bacteriophage engineering"

**Table S1.** List of the primers employed in this study for the amplification of both plasmid backbones and insert sequences.

| Plasmid name | Backbone | Insert | Function | Cloning primers | Primer Sequence (5'-3') |
|---|---|---|---|---|---|
| $\text{pEdit}_{\text{Early}}$ | pLEB579 | $nluc_{\text{Early}}$ | Homology donor template for engineering of phage K::$nluc_{\text{Early}}$ | pLEB579_Fw | AGTCGATGTTAAACCGTGTGCTCTACG |
| | | | | pLEB579_Rev | CGCGCTATTAATCGCAACATCAAACC |
| | | | | Gene_String_Fw | GGGGCTTTTATTTTGGTTTGATGTTG |
| | | | | Gene_String_Rev | TTATAGTTTTGGTCGTAGAGCACACG |
| $\text{pEdit}_{\text{Middle}}$ | pLEB579 | $nluc_{\text{Middle}}$ | Homology donor template for engineering of phage K::$nluc_{\text{Middle}}$ | pLEB579_Fw | AGTCGATGTTAAACCGTGTGCTCTACG |
| | | | | pLEB579_Rev | CGCGCTATTAATCGCAACATCAAACC |
| | | | | Gene_String_Fw | GGGGCTTTTATTTTGGTTTGATGTTG |
| | | | | Gene_String_Rev | TTATAGTTTTGGTCGTAGAGCACACG |
| pEdit_Late | pLEB579 | $nluc_{\text{Late}}$ | Homology donor template for engineering of phage K::$nluc_{\text{Late}}$ | pLEB579_Fw | AGTCGATGTTAAACCGTGTGCTCTACG |
| | | | | pLEB579_Rev | CGCGCTATTAATCGCAACATCAAACC |
| | | | | Gene_String_Fw | GGGGCTTTTATTTTGGTTTGATGTTG |
| | | | | Gene_String_Rev | TTATAGTTTTGGTCGTAGAGCACACG |
