## Supplemental Table 2 for "Computational pipeline for targeted integration and variable payload expression for bacteriophage engineering"

**Table S2.** Synthetic DNA sequences employed as homology donors to facilitate targeted homologous recombination.

| Gene String Name | Length (bp) | Sequence (5'-3') |
|---|---|---|
| nluc_Early | 1317 | AAAGTCGAAGGGGGCTTTTATTTTGGTTTGATGTTGCGATTAATAGCGCGAGAAGGTTATAATAAAGAATACTTGTCTAAAGCACTCAGATTAATCAATGACCATGCTCCTAGGGAGTTAAGTTATGATTTTAATAATGTAGAAGCGGATGTTAATATTCACACAATGTTATATGTTAAACCTGAAGATAGATTTATATATAAGGATATATCCTATGACTTCCCGGGTGATTTAATTATTTGTATAGTTGATGATGATGCTATTGTATACCATCAAGGTGAGCAGATTTCAGGTATTAGTATTTTAAGAATACTAGAAGAGATATTTTAAGAGGAGGTAAATATATATGGTATTCACTTTAGAAGATTTCGTAGGTGATTGGCGTCAAACTGCTGGTTACAACTTAGATCAAGTATTAGAACAAGGTGGTGTATCATCATTATTCCAAAACTTAGGTGTATCAGTAACTCCAATCCAACGTATCGTATTATCAGGTGAAAACGGTTTAAAAATCGATATCCACGTAATCATCCCATACGAAGGTTTATCAGGTGATCAAATGGGTCAAATCGAAAAAATCTTCAAAGTAGTATACCCAGTAGATGATCACCACTTCAAAGTAATCTTACACTACGGTACTTTAGTAATCGATGGTGTAACTCCAAACATGATCGATTACTTCGGTCGTCCATACGAAGGTATCGCTGTATTCGATGGTAAAAAAATCACTGTAACTGGTACTTTATGGAACGGTAACAAAATCATCGATGAACGTTTAATCAACCCAGATGGTTCATTATTATTCCGTGTAACTATCAACGGTGTAACTGGTTGGCGTTTATGTGAACGTATCTTAGCTTAATTTAAGGAGGATAAGTAATCATGATAGGAATAACAATATTAATTACGATAATGAGTATATCAACTATCTCTATGTATATTTATTTTTTAGTAGACTTGATTCAGTCAATCAGATATAATAGTTTTGATAAAGTAATTAACGTCATAACATTTGTACTTATGACAGTTATAATAGCATCAGGTATTTTAGCTATACTTGGAATATAGAGCTCATTTAAGAAGCGGTTAAGTAGTTAGAGGGGATTTGTCCTAAAATAGTATACCGCTTCTATATGGAAGGCTGAGAGGTCTTAGAATTGAAAGGAGAGATATAATGATTCATATATTTTTAACTGATAGTTATGATAATAAAGTTTTAAATACTGTACTCAGATATATTAATACTACTAGTGATAGAGAGCTTAGTAGTCGATGTTAAACCGTGTGCTCTACGACCAAAACTATAAAACCTTTAAG |
| nluc_Middle | 1435 | AAAGTCGAAGGGGGCTTTTATTTTGGTTTGATGTTGCGATTAATAGCGCGGCAACTTTCTCCATTCGAACAATCAAAACATCACACACAAACAATTTATGATGATATTCAAGTACTAGATATGATTATTTCTAAAGGTGCAAAAGGATTAGAGTTTGTGGAAACTTTAGACCCTGCTTTAATGATACGTGCAATGGAAACTAAAGATAAGATTACCGGAAATCAATTAAAAGGTATGTCATTTATTGGACTTAGAGAATTACAATTAAAACAAACAGCTCAAGATACAGCTATGAGTGAAGTATTATTAGAATTTATACCTGAAGAGAAACATGAAGAGGTATTACAACGATTAGAAGAACTACAAAATGAATTCTACAAAAATCTAGATTTAGATGAAGAAAGTAGAAAATTAAAAGAAGCTCTTGATAGAGTAGGCTATACAATTTAGGAGGAGGTAAATATATATGGTATTCACTTTAGAAGATTTCGTAGGTGATTGGCGTCAAACTGCTGGTTACAACTTAGATCAAGTATTAGAACAAGGTGGTGTATCATCATTATTCCAAAACTTAGGTGTATCAGTAACTCCAATCCAACGTATCGTATTATCAGGTGAAAACGGTTTAAAAATCGATATCCACGTAATCATCCCATACGAAGGTTTATCAGGTGATCAAATGGGTCAAATCGAAAAAATCTTCAAAGTAGTATACCCAGTAGATGATCACCACTTCAAAGTAATCTTACACTACGGTACTTTAGTAATCGATGGTGTAACTCCAAACATGATCGATTACTTCGGTCGTCCATACGAAGGTATCGCTGTATTCGATGGTAAAAAAATCACTGTAACTGGTACTTTATGGAACGGTAACAAAATCATCGATGAACGTTTAATCAACCCAGATGGTTCATTATTATTCCGTGTAACTATCAACGGTGTAACTGGTTGGCGTTTATGTGAACGTATCTTAGCTTAATAGATAGTGAGGTTAGAGTAATGGCAGATGAGATTAGTTTAAATCCAATACAAGATGCTAAGCCAATTGACGATATAGTAGATATCATGACATACTTAAAAAACGGGAAAGTACTGAGAGTTAAACAAGACAATCAAGGAGATATCCTTGTTAGAATGAGTCCAGGGAAACACAAATTTACTGAAGTATCTAGAGACTTAGATAAAGAATCATTCTACTATAAAGGCATTGGGTTCTCTATAATGTATCTGTTAACTCTCTTATAACATTTGATGTTTATCTAGATGAAGAATATTCAGAAACAACTAAGGTTAAGTATCCTAAAGATACTATTGTAGAATATACAAGAGAAGACCAAGAAAAAGATGTTGCTATGATTAAAGAAATACTTACAGATAATAAAGTCGATGTTAAACCGTGTGCTCTACGACCAAAACTATAAAACCTTTAAG |
| nluc_Late | 1382 | AAAGTCGAAGGGGGCTTTTATTTTGGTTTGATGTTGCGATTAATAGCGCGGTGTTTTACATAAATGAAATCTGAGAATTCATATGTGCGGTTGACGAACCGGTTTGCTGTGTCCTCGAATGAGGGTAGGATTTCATTGAGTATATAGCGCAAGTGTTTGTTCCTAGGATACGATACTAGCTATATACTTTACATATTTAAAAGGAGAGATACAAATGGCATCAGCAAAACAATTATATTATACGGAAAGTTTAGTTGGTAAGGCAATTATTAATAATAAAGTATCTAATAAAGAAGAAGTTTGGGATAAGTTAGAGCTACTTCCTGAAACTAAATTAGAAGATTTAGATAACAAACAAATGTCTGAAGTTATCAAAAAACTAAATCAAATTAATGAGTAAGAGGAGGTAAATATATATGGTATTCACTTTAGAAGATTTCGTAGGTGATTGGCGTCAAACTGCTGGTTACAACTTAGATCAAGTATTAGAACAAGGTGGTGTATCATCATTATTCCAAAACTTAGGTGTATCAGTAACTCCAATCCAACGTATCGTATTATCAGGTGAAAACGGTTTAAAAATCGATATCCACGTAATCATCCCATACGAAGGTTTATCAGGTGATCAAATGGGTCAAATCGAAAAAATCTTCAAAGTAGTATACCCAGTAGATGATCACCACTTCAAAGTAATCTTACACTACGGTACTTTAGTAATCGATGGTGTAACTCCAAACATGATCGATTACTTCGGTCGTCCATACGAAGGTATCGCTGTATTCGATGGTAAAAAAATCACTGTAACTGGTACTTTATGGAACGGTAACAAAATCATCGATGAACGTTTAATCAACCCAGATGGTTCATTATTATTCCGTGTAACTATCAACGGTGTAACTGGTTGGCGTTTATGTGAACGTATCTTAGCTTAAGTGTTAAATTAATATAGATAAACATGAGCGACCTACTGTTATATTATTGTTAGAAATAAATATAATAGAAAGGTCGGTTTTTTAATGGCTAATGAAACTAAACAACCTAAAGTTGTTGGAGGAATAAACCTTAGCACAAGAACTAAGAGCAAAACATTTTGGGTAGCAATTATATCAGCAGTAGCATTATTTGCTAACCAAATTATAGGTGCTTTCGGTTTAGACTACTCAGCTCAAATTGAGCAAGGTGTAAATATTGTAGGTTCTATACTAACACTATTAGCAGGTTTAGGTATTATTGTTGATAATAATACTAAAGGTCTTAAAGATAGTGATATTGTTCAAACAGACTATCTTAAACCTCGTGATAGTAAAGACCCTAATGAATTCGTTCAATGGCAGTCGATGTTAAACCGTGTGCTCTACGACCAAAACTATAAAACCTTTAAG |
